# Maize RAMOSA3 accumulates in nuclear condensates enriched in RNA POLYMERASE II isoforms during the establishment of axillary meristem determinacy

**DOI:** 10.1101/2021.04.06.438639

**Authors:** Edgar Demesa-Arevalo, Maria Jazmin Abraham-Juarez, Xiaosa Xu, Madelaine Bartlett, David Jackson

## Abstract

Maize meristem determinacy is regulated by Trehalose-6-Phosphate Phosphatase (TPP) metabolic enzymes RAMOSA3 and TPP4. However, this function is independent of their enzymatic activity, suggesting they have an unpredicted, or moonlighting function. Using whole-mount double immunolabeling and imaging processing, we investigated the co-localization of RA3 nuclear speckles with markers for transcription, chromatin state and splicing. We find evidence for RA3 co-localization with RNA POL II, a transcription marker, and not with markers for promoter chromatin remodeling or mRNA processing, suggesting a function of nuclear RA3 in mediating a transcriptional response during meristem determinacy.

## Introduction

Meristems are niches of undifferentiated stem cells that coordinately specify new organs during the plant life cycle. Maintenance of meristem activity determines plant architecture, including the seed bearing inflorescences^1^. Maize is monoecious, however male (the tassel) and female (the ear) inflorescences follow similar initial development, generating axillary meristems known as spikelet pair meristems (SPM) from the inflorescence meristem (IM). In tassels, branch meristems give rise to the branched spike, whereas ear IMs are more determinate, and only generate SPMs. This determinacy is lost in *ramosa* mutants, generating branch meristems from early SPMs in the ear, and branched cobs^2,3^. Maize has three *ramosa* mutants, *RAMOSA1* (*RA1*) and *RA2* encode C2H2-zinc finger^4^ and LATERAL ORGAN BOUNDARY^5^ transcription factors respectively. Interestingly, RA3 encodes a trehalose-6-phosphate phosphatase (TPP), an enzyme responsible for catalysis of trehalose from trehalose-6-phosphate (T6P), suggesting that trehalose or its precursor trehalose-6-phosphate (T6P) has the potential to promote differentiation^6^. T6P has been proposed as a putative signal involved in many biological processes^7^.

In Arabidopsis, mutants in TREHALOSE-6-PHOSPHATE SYNTHASE 1 (TPS1), the enzyme catalyzing T6P synthesis, are embryo lethal^8^, but can be rescued by expressing TPS1 using inducible or tissue specific promoters during seed development, however this leads to a delay in flowering. In contrast, the misexpression of TPS1 in meristematic tissues accelerates flowering, suggesting T6P plays a crucial role in this process^9^. It is proposed that T6P acts as a signal and negative feedback regulator of sucrose levels, linking sugar resources to growth and development. This feedback may be integrated by the sucrose non-fermenting-1 related protein kinase 1 (SnRK1), since T6P is able to inhibit its kinase activity, however the molecular mechanisms by which T6P or trehalose are involved remain unknown^10,11^.

T6P regulates branching in maize^6^ and Arabidopsis^12^ and a RA3 paralog, TPP4, acts redundantly to suppress branching in maize ears, since *ra3; tpp4* double mutants have enhanced branching. Intriguingly, this phenotype is independent of their catalytic activity, because a catalytically inactive version of RA3 expressed under its native regulatory elements can partially complement the *ra3* phenotype. In support of this hypothesis, *tpp4* weak alleles with ~35% of wild type activity have the same quantitative effect in enhancing *ra3* as null alleles. Together, these observations suggest that RA3 and TPP4 suppress branching independently of their enzymatic function. RA3 and TPP4 localize to the cytoplasm and nuclei when are expressed transiently in tobacco, and a native antibody against RA3 found its localization in cytoplasmic and nuclear speckles in cells subtending the SPMs in developing tassels and ears. This localization pattern supports a moonlighting hypothesis, as reported for other sugar metabolic enzymes that also localize in nuclei^13^.

Nuclear speckles, historically defined as a punctate nuclear localization pattern, are compartmentalized condensates with a dense accumulation of proteins and RNA observed in transmission electron micrographs of nuclei. They were later identified as ribonuclear aggregates enriched in splicing factors and/or transcriptional machinery^14^. Nuclear condensates are related to many biological processes, and it was recently demonstrated that different transcription stages are segregated into different speckles, and their compartmentalization is established by the phosphorylation status of the Carboxy Terminal Domain of RNA POLYMERASE II (POLII-CTD)^15^. Current models of chromatin organization also point toward molecular condensation in aggregates mediated by histone marks and protein interactions^16^, suggesting many nuclear processes are organized in membrane-less structures^17^. All transcription stages, from chromatin decondensation, transcriptional elongation and mRNA processing, are dynamically compartmentalized in different speckles^18^. Here we asked if nuclear speckles containing RA3 are related to chromatin, splicing or transcriptional processes, by adapting a whole-mount protocol to perform double immunolabelling in developing ears. We used a native antiserum against RA3, and a range of nuclear markers associated with nuclear speckles to estimate 3D colocalization and find evidence for preferential co-localization of RA3 with RNA pol II, suggesting it acts in transcriptional control.

## Results

We developed a whole mount protocol using thick vibratome sections, modified from (^19^) for immunolocalization in 2-5 mm maize ear primordia, where the first molecular and phenotypic cues associated with branching are distinguishable^20^. This stage is enriched in growing SPMs and has high RA3 expression. To verify this protocol was consistent with previous results using semi-thin sections^13^, we confirmed RA3 localization in cytoplasmic and nuclear speckles in the cells subtending developing SPMs in wild type (B73) (**Figure 1A** **and** **C**). We used *ra3* mutant ears as a negative control to verify antibody specificity, as suggested in similar protocols^21^, and as expected no signal was detected (**Figure 1B**). Since RA3 expression was found in a limited number of cells, we used a positive control, anti-YSPTSPS repeat from the C-terminal domain of RNA polymerase II (POL II-CTD) phosphorylated on serine 2 (Ser2 Pho) as a constitutive nuclear marker, and found uniform nuclear immunostaining throughout the whole inflorescence, including in cells where RA3 was expressed, corroborating the technical consistency in our immunolabeling protocol **(Figure 1A-B)**.

**Figure 1.**
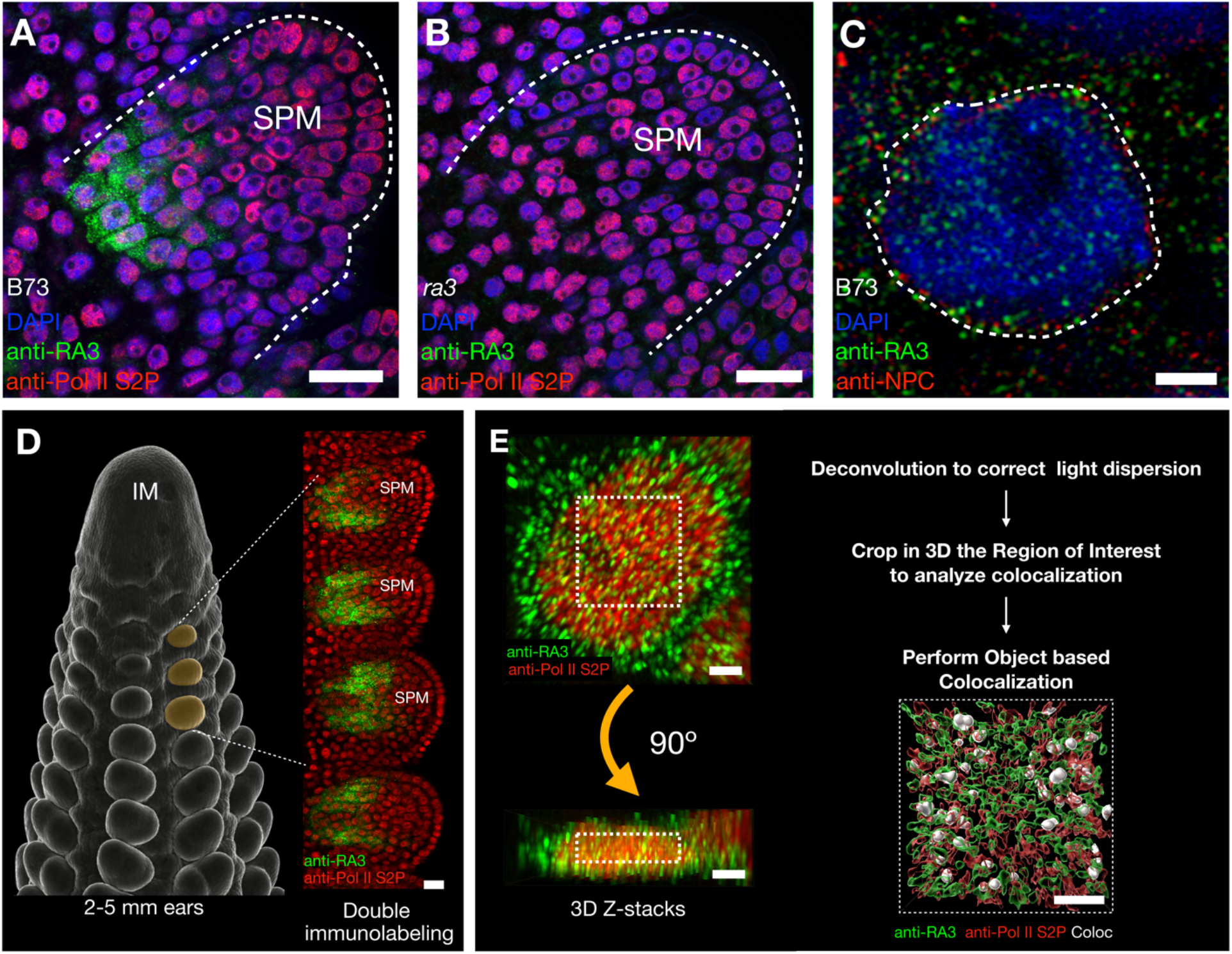
RA3 localization and workflow for co-localization in nuclear speckles during maize spikelet pair determinacy. **A)** Representative images of whole-mount immunolocalization in maize inflorescences showing the expected pattern for RA3 (green specles) subtending the base of SPMs. **B)** *ra3* mutant was used as a negative control, showing the absence of signal **C)** RA3 (green) is localized in nuclear and cytoplasmic speckles, the nuclear envelope was labeled using an anti-Nuclear Pore Complex protein antibody (anti-NPC, in red), the nucleus shape is highlighted by the white dashed line, a representative median confocal section shows the nucleus surrounded by the NPC, with RA3 nuclear speckles inside. Sections were counterstained with DAPI (blue). **D)** Axillary spikelet pair meristem (SPM) development in ears occurs continuously on the flanks of the Inflorescence Meristem (IM), image on left is a scanning electron micrograph; whole-mount double immunolocalizations were performed using 2-5 mm ears. 3D Z-stacks of nuclei were acquired from the uppermost three SPMs (marked orange on the SEM) in cells expressing RA3 and covering entire nuclei (~5 μm). **E)** after acquisition, images were processed by deconvolution and cropping a 3D region of interest inside the nucleus (dashed lines) and the percentage colocalization was calculated by volume (object based) and rendered for easy representation and visualization. Scale bars, **A-C and E**: 2 μm, **D**: 20 μm.

Nuclei were imaged from cells subtending SPMs prior to their elongation, typically the uppermost 3 developing SPMs^20^ prior to their division into SMs, and the stage where the potential to become a branch is repressed (**Figure 1D**). Images were acquired as Z-stacks of single nuclei, followed by deconvolution and background subtraction. Next, we visualized RA3 distribution in 3-D, where we cropped the cuboid volume inside each nucleus that was used to calculate the object-based colocalization between RA3 and the different nuclear markers, expressed as a percentage of volume (**Figure 1E**). To verify the reproducibility of our results, we performed at least two independent experiments, each using ears from at least 5 plants, and imaged at least 14 nuclei per marker or controls.

Since RA3 speckles were also observed throughout the cytoplasm, we first sought to demonstrate that they were indeed inside nuclei, and not in the cytoplasm above or below. For this, we performed double immunolabeling using an anti-Nuclear Pore Complex protein (a-NPC) antibody. This outlined the nuclear envelope and allowed us to image medial sections of individual nuclei to distinguish RA3 speckles inside of the nucleus (**Figure 1C and Supplementary Figure 1A**). a-NPC was detected using a different secondary antibody with a different fluorophore tag, allowing us to distinguish its localization from that of RA3, and in some experiments we also imaged nuclei using 4′,6-diamidino-2-phenylindole (DAPI). We imaged each marker using appropriate lasers and acquiring a narrow range of emission (~30 nm) on the confocal microscope to avoid cross-talk. To better visualize RA3 inside nuclei, we performed Z-Stack acquisition, 3D reconstruction, and rendering of individual nuclei. Using DAPI as a reference we could clearly visualize RA3 speckles inside the nuclei and wrapped by the NPC proteins in the nuclear envelope (**Supplementary Figure 1B-C**).

We next asked if we could identify the sub-nuclear compartment(s) or condensates in which RA3 was localized, by performing double immunolabelling using antibodies against nuclear markers, followed by image processing to quantify the percentage of volumetric colocalization between RA3 and selected markers ^22–24^. The transcription cycle can be divided in five main steps: chromatin remodeling, preinitiation, initiation and pausing, active elongation and mRNA processing^25^ (**Figure 2A**). Chromatin remodeling in promoters can be monitored using histone marks associated with accessible or compact chromatin. We selected Histone H3 acetylation in Lysine 9 (H3K9Ac) as a marker for euchromatin, since this mark is associated with open promoters and enhancers in maize^26^, and two histone modifications associated with heterochromatin; H3K9me2^27^ for constitutive heterochromatin and H3K27me3 for facultative heterochromatin^28^. We found that RA3 was significantly enriched in co-localization with H3K9Ac (16.6%, n=30) compared to H3K9me2 (13.2%, n=33) or H3K27me3 (12.0%, n=27) (**Figure 2B-D, M**). We confirmed the specificity of the heterochromatin markers by negative controls, showing their reduced colocalization with RNA POL II as a marker for active transcriptional elongation, labelled by the anti-YSPTSPS repeat Ser2 Pho from POL II-CTD (H3K9me2/ Ser2 Pho POL II-CTD =8.9%, n= 20; H3K27me3 = Ser2 Pho POL II-CTD 8.2%, n= 14) (**Figure 2K-L, M**). Our results suggest that RA3 speckles preferentially co-localized with euchromatin related condensates. These estimates using confocal microscopy Z-stacks and image processing were verified using Structured Illumination Microscopy using a DeltaVision OMX super-resolution microscope, and reconstructed entire nuclei in double immunolabelling of RA3 and H3K9me2. We estimated ~14.3% (n=3) colocalization which was not significantly different (*P-value* = 0.4, Mann-Whitney test) to our estimates using confocal acquired images (**Supplementary Figure 3**).

**Figure 2.**
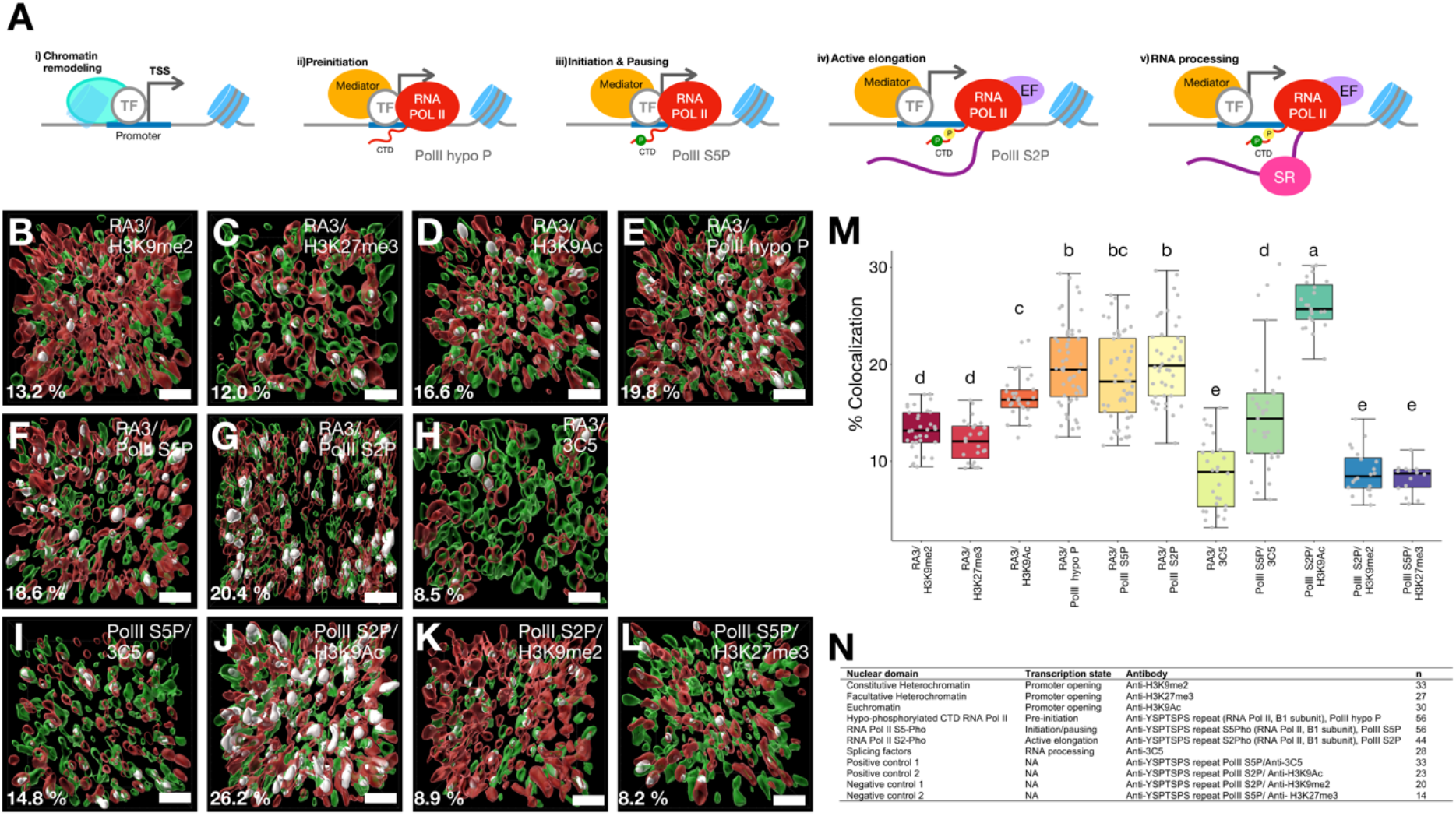
RAMOSA3 accumulates in nuclear condensates enriched in RNA POLYMERASE II. **A)** The transcription process can be described by five key processes i) chromatin remodeling to open promoters; ii) pre-initiation, when RNA POL II binds to promoters, and is characterized by its hypo-phosphorylated CTD (red tail); iii) initiation, where the POLII CTD is phosphorylated on Ser5 (green circle), and if the gene is regulated by pausing the CTD maintains only S5Pho, but if it is transcribed directly the CTD is also phosphorylated on Ser2 (iv), yellow circle) to initiate active elongation; v) splicing is coupled to the transcription process and can be identified by serine/arginine-rich SR proteins (magenta). TF: Transcription Factor, EF: Elongation Factors. **B-H)** Representative renderings of RA3 (green) and transcription associated markers (red), colocalization is highlighted in white, the percentage shows the average of % of RA3 colocalization volume with respect to the marker. **B)** H3K9me2 and **C)** H3K27me3 mark heterochromatin; **D)** H3K9Ac marks euchromatin; **E)** anti-RNA POL II CTD repeat YSPTSPS non-phosphorylated marks pre-initiation; **F)** anti-YSPTSPS repeat Ser5 phosphorylated marks initiation and pausing; **G)** anti-YSPTSPS repeat Ser2 phosphorylated marks active elongation; **H)** splicing condensates, using the 3C5 antibody marks SR proteins. Controls for specificity were **I)** positive control RNA POL II CTD repeat YSPTSPS versus 3C5, **J)** positive control anti-YSPTSPS repeat Ser2 phosphorylated H3K9Ac, **K)** negative control anti-YSPTSPS repeat Ser2 phosphorylated versus H3K9me2, and **L)** negative control anti-YSPTSPS repeat Ser2 phosphorylated versus H3K27me3. **M)** the colocalization percentages were compared by a Kruskal-Wallis test (*P-value* < 0.001) and statistical groups were determined by Dunn-Bonferroni multiple comparison, each dot represent the percentage for individual nuclei. **N)**Antibodies used for specific transcriptional stages and number of nuclei analyzed per each marker, all data come from at least 2 independent experiments. Scale bars: 1 μm.

To further analyze the possibility that RA3 preferentially co-localized with transcriptionally active chromatin, we characterized different transcription stages by colocalization with stage-specific markers. First, we used an antibody against the hypo-phosphorylated YSPTSPS repeat from RNA POL II-CTD, which is characteristic, together with Mediator, for transcriptional pre-initiation^29,30^. RA3 colocalized with hypo-phosporylated POL II (PolII hypo P) at 19.8% (n= 56, **Figure 2E, M**). To analyze if RA3 was more enriched during initiation and pausing, or during active elongation, we next used anti-YSPTSPS repeat Ser5 Pho (PolII S5P) or anti-YSPTSPS repeat Ser2 from POL II-CTD (PolII S2P) which label modifications present in those stages, respectively^25,31,32^. RA3 showed similar colocalization with the active elongation marker (20.4%, n=56) and with the initiation and pausing marker (18.6%, n=44) (**Figure 2F-G, M**). Co-localization of RA3 with the different transcriptional stage markers was consistently higher than with the heterochromatin markers and was higher though not significant than with the euchromatin marker, suggesting RA3 is involved in active transcriptional elongation.

We next asked if RA3 speckles co-localized with RNA splicing, another nuclear process associated with condensates. We measured colocalization with serine/arginine-rich (SR) proteins, RNA binding proteins that perform splicing, using the 3C5 monoclonal antibody^33^. In agreement with previously reported immunolocalization experiments using semi-thin sections^13^, RA3 colocalization with 3C5 was relatively low (8.5%, n=28, **Figure 2H, M**) and not significantly different from negative controls, confirming that RA3 nuclear speckles were not associated with post-transcriptional RNA processing. Representative images before processing for all markers are shown in **Supplemental Figure 2**. In summary, RA3 accumulated in the nuclei of cells subtending the SPM in nuclear speckles that co-localized with markers for euchromatin and different stages of the RNA POL II transcription cycle marked by its CTD phosphorylation state, but not with markers for heterochromatin or mRNA processing. Our results suggest that RA3 could be mediating a specific transcriptional program, for example by guiding the interaction of transcription or elongation factors with the RNA POL II complex. This may activate genes involved in establishing the determinate state of SPMs in response to developmental signals, such as T6P.

## Discussion

Complementation of *tps1-1* null-alleles in arabidopsis using *E. coli* TPS (OtsA) restores the endogenous levels of T6P and complements embryo lethality and branching, suggesting T6P rather than trehalose itself is a developmental signal^12^. T6P has also been postulated as a flowering signal^34,35^. T6P is a substrate of TPP in trehalose biosynthesis, however, loss or reduction of RA3 or TPP4 enzymatic activity in maize are not correlated with degrees of branching, suggesting RA3 has a moonlighting function, which could involve T6P binding and/or signaling T6P levels. Similar moonlighting roles have been proposed for nuclear HEXOKINASE1 in *A. thaliana*^36^ and other metabolic enzymes and their substrates^37,38^. Interestingly, GFP-TPS1^9^ and TPPs in arabidopsis^39^ or expressed heterologously in *Nicotiana*^40^ are also nuclear localized.

Using whole-mount immunolabeling and quantitative 3D colocalization, we found that RA3 accumulated in cytoplasmic and nuclear speckles. The identity of the RA3 cytoplasmic speckles is unknown, however for the nuclear speckles we observed highest co-localization with RNA POL II, either in pre-initiation, initiation and active elongation stages of the transcription cycle. In contrast, we observed much lower co-localization, close to background levels, with markers for heterochromatin, chromatin remodeling or mRNA splicing. Analysis of the transcription cycle in animal cells by immunolocalization and 3D colocalization found from ~8% to ~34% between RNA POL II and other markers, similar to our findings^15^. Current models for transcription suggest a compartmentalization of condensates that are dynamic in their composition and function, and exchange components rather than forming new speckles. Our results suggest that RA3 could mediate a specific developmental response by interacting with specific components of the transcriptional machinery during the establishment of SPM fate and promotion of branch determinacy (**Figure 3**). RA3 may be localized in nuclear speckles by a liquid-liquid phase separation mechanism, since it contains a predicted nuclear localization signal^41^ and four putative disordered or low complexity regions, which are associated with proteins that form condensates by this mechanism^42^, however this remains to be confirmed experimentally (**Supplemental Figure 4**). Interestingly, when RA3 is expressed heterologously in *Nicotiana,* it shows a non-speckled distribution in both cytoplasm and nucleus, suggesting the ability to form speckles or condensates may be dependent on additional factors present in the endogenous RA3 expression domain^13^.

**Figure 3.**
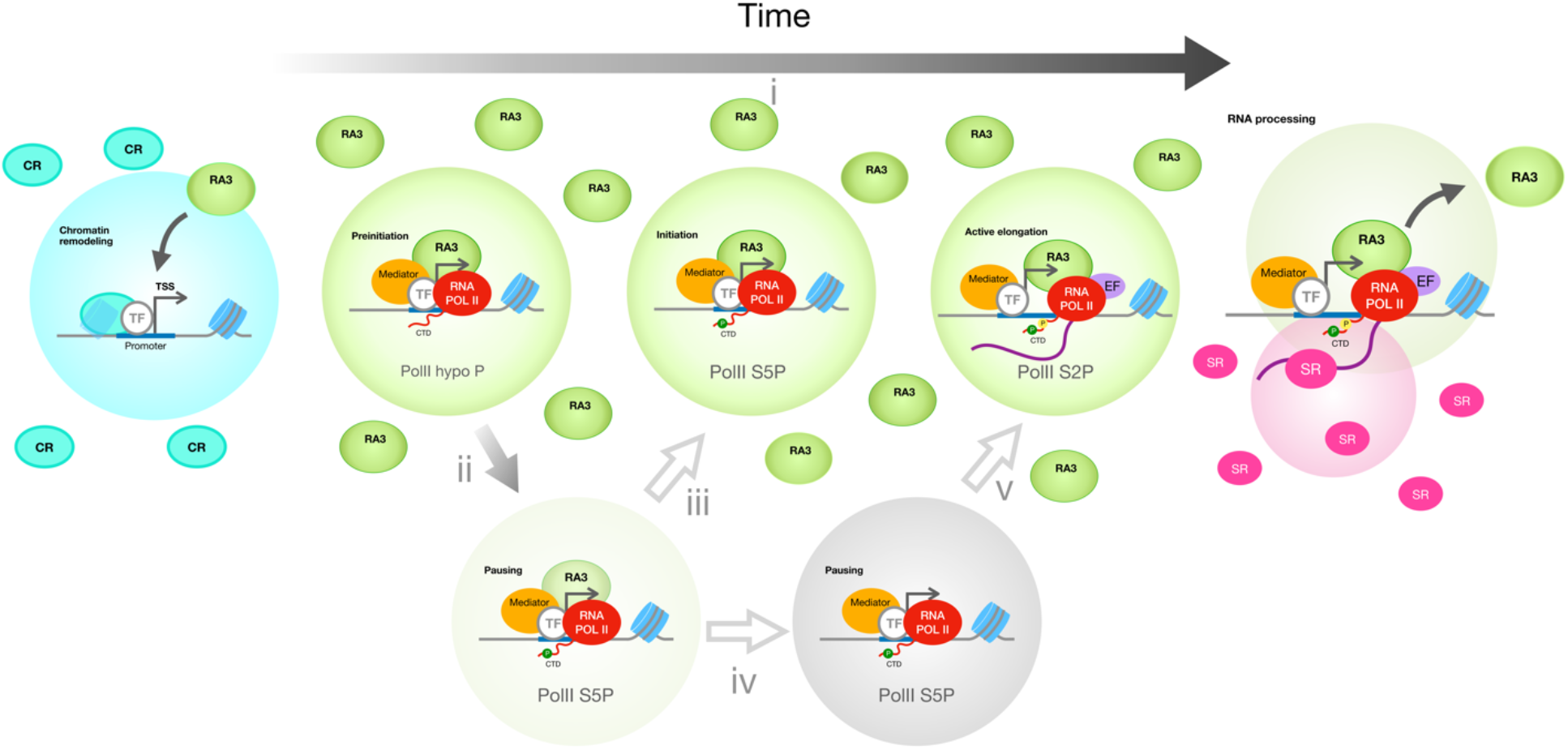
A model depicting the enrichment of RA3 in different stages of transcription. **i)** Prior to initiation the chromatin remodelers (CR) and histone modifier enzymes establish the status of promoters; an open status allows Transcription Factors (TF) to bind promoter *cis* elements. During pre-initiation, RA3 is enriched in condensates with hypophosphorylated CTD RNA POL II. Phosphorylation of the CTD in Ser5 (green circle) leads to initiation, however this phosphorylation is also present in loci subject to pausing regulation. RA3 also colocalized in CTD-Ser5p containing condensates and with those accumulating the hypophosphorylated CTD in RNA POL II or with the CTD-Ser2p (yellow circle) specific for transcriptional elongation, EF: Elongation Factors.

## Methods

### Whole-mount immunolocalization

The antibody against RA3 used in this study was generated as was described previously^13^. Whole mount immunolocalization was performed using 2-5 mm ears collected from wild type B73 and *ra3* (in B73 background) plants around 6 weeks after sowing. The ears were immediately pre-fixed in fixative solution, 4% paraformaldehyde, 2% Tween-20 in Phosphate Buffered Saline (PBS), for one hour at 4ºC. After rinsing with PBS, the ears were embedded in 6% Agarose in PBS and sectioned at 100 μm using a vibratome (Leica) and fixed for 2 hours more at 4°C with gentle agitation. The sections were washed three times with PBS and then digested using a cell wall digestion enzyme solution containing: 1% Driselase (Sigma-Aldrich), 0.5% cellulase (Sigma-Aldrich), 0.75% Pectolyase Y-23 (Duchefa Biochemie) for 12 minutes at room temperature. The tissue was rinsed three times with PBS and permeabilized for 2 h in PBS, 2% Tween-20, rinsed 2 times with PBS and blocked with 4% Bovine Serum Albumin (BSA, Sigma) for 1 h, and then incubated with the primary antibodies overnight at 4°C. The samples were washed for 8h at 4°C with gentle agitation using 0.2% Tween-20 in 1X PBS, replacing it every 2 hours, and then incubated with the secondary antibodies overnight at 4°C. The tissue was washed for 6 h with 0.2% Tween-20 in 1X PBS replacing the solution every 2 hours. Tissues were counterstained with DAPI (4′,6-diamidino-2-phenylindole, 1 μg/ml in PBS) and mounted with ProLong Gold (ThermoFisher Scientific) using #1.5 coverslips (thickness: 0.17 mm) and sealed with nail polish. The antibodies and the concentrations used in this study can be found in **Supplemental Table 1**. All immunolocalizations were carried out at least 2 times to verify biological and technical consistency.

### Imaging double labelling immunolocalizations

All images used for quantification of colocalization were acquired using a Zeiss LSM 780 confocal microscope using a 63X objective, performing multi-tracking scanning in each channel independently using photon-counting mode with the appropriate filters (~20 nm of emission collected) per fluorophore to minimize cross-talk between fluorophores. To image entire nuclei (~3 μm diameter), the nuclear markers were used as a reference for Z-stacks and were captured at minimum speed and with the pinhole aperture calibrated to cover the same optical slice in every track with the optimal resolution and step-size. Additional images, including anti-NPC, were acquired using a Zeiss LSM 900 using the Airyscan 2 mode under equivalent parameters, or using 3D Structured Illumination Microscopy on a DeltaVision OMX super-resolution microscope under a 100X objective.

### Image processing for Colocalization quantification

3D Z-stacks taken by the LSM780 were processed using IMARIS 9.5.1 software, first correcting light dispersion using the deconvolution module with the required parameters (magnification: 63X Numerical Aperture 1.4; Oil diffraction index:1.14; mounting medium: ProLong Gold, 1.47 Refraction Index) using 10 iterations. The deconvoluted images were background subtracted using standard thresholding and the filtered images were used to calculate object-based colocalization by the colocalization module in IMARIS 9.5.1. The processed images then were used to render the 3D models to visualize the data.

### Statistical analysis

Data plotting and statistical analysis were performed using R Studio Version 1.2.5033. The percentages of volume colocalization were compared using a Kruskal-Wallis test and statistics groups were determined by Dunn-Bonferroni multiple comparison. Micrographs and 3D rendering models were compiled using ImageJ 2.1.0/1.53c, IMARIS Viewer 9.5.1 and Keynote 10.3.5.

## Acknowledgements

We thank Daniel Grimanelli for his technical advice adapting the protocol for immunolabeling. Nour El-Amine and Tse-Luen Wee from the CSHL Microscope Shared facility for their technical advice operating the OMX Super-resolution microscope and using IMARIS for colocalization analysis, David Spector for providing the 3C5 antibody and Tim Mulligan and Kyle Schlecht for plant care. This project was supported by funding from the National Science Foundation (IOS-1755141), and the National Council of Science and Technology (CONACYT-Mexico to E. D.-A.)

## Authors contributions

E.D.-A., designed and performed all experiments and analysis, prepared figures and co-wrote the manuscript. M.J.A.-J and X.X. purified the RA3 antisera under the supervision of M.B. D.J. supervised the research and co-wrote the manuscript.

---

**Supplemental Figure 1.**
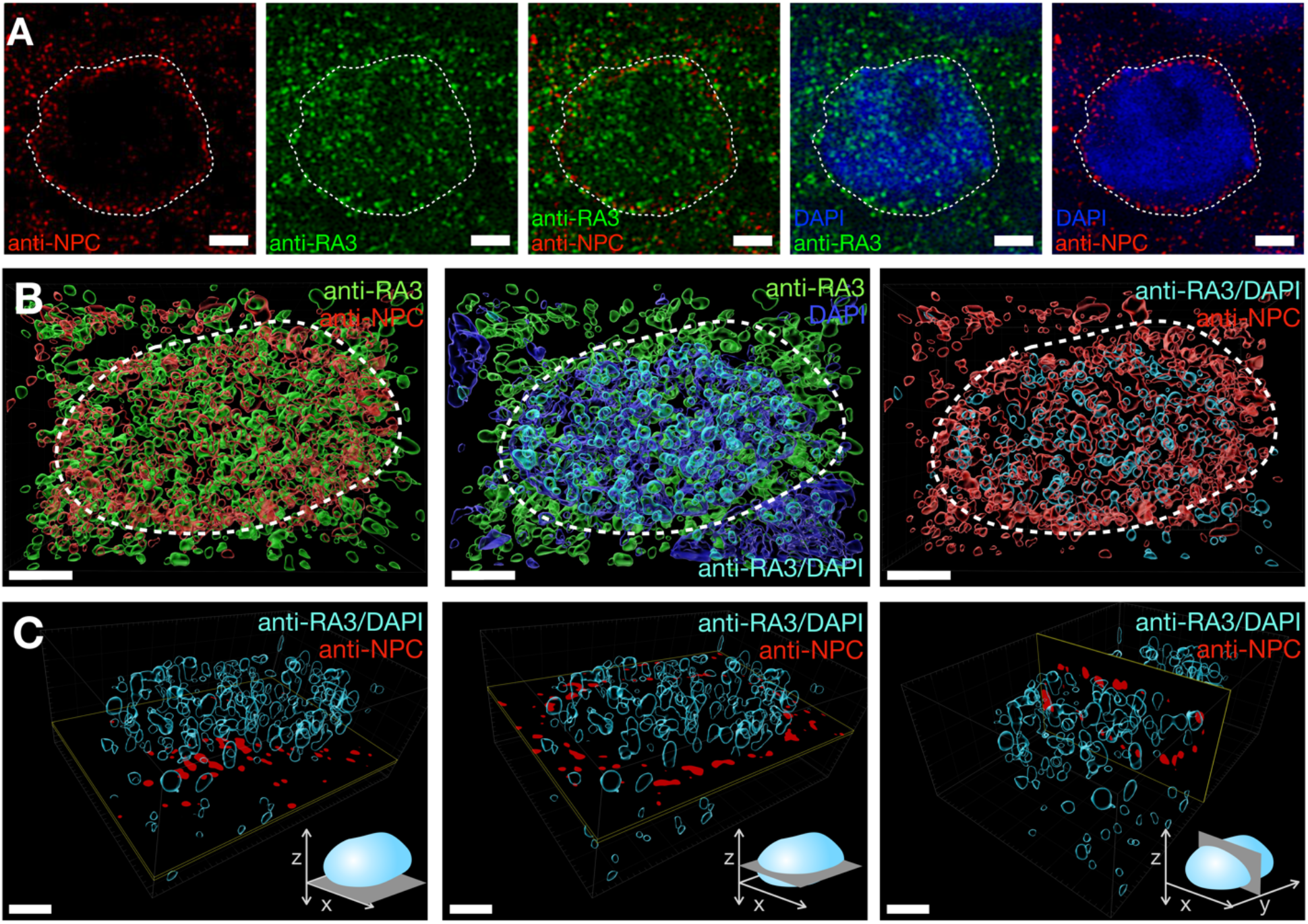
3D rendering of RA3 and Nuclear pore complex proteins (NPC) confirms RA3 localization inside the nucleus. **A)** Representative images of confocal micrographs labeling NPC (red) and RA3 (green), showing RA3 speckles inside the nucleus (white dashed line), overlapping RA3 and NPC or nucleus staining with DAPI corroborate nuclear localization of RA3. **B)** Representative rendering of one Z-stack immunolabelling a nucleus with anti-RA3 (green) and anti-NPC (red), next pictures in the same panel are showing the counterstaining with DAPI (blue) and RA3 speckles colocalizing with DAPI (cyan) to identify the nuclear RA3 and rendering of NPC and nuclear RA3 in the entire Z-stack. **C)** For better visualization of nuclear RA3, a transverse section (XY plane) of NPC signal was placed at the bottom of the nucleus (left panel), or in the middle of the nucleus (middle panel), NPC clearly show the signal around nuclear RA3. Sagittal (XZ plane) section (right panel) in the middle of the same nucleus also shows the NPC signal surrounding nuclear RA3, the drawings in **C)** show the section position in each reconstruction. Scale bars: 2 μm.

**Supplemental Figure 2.**
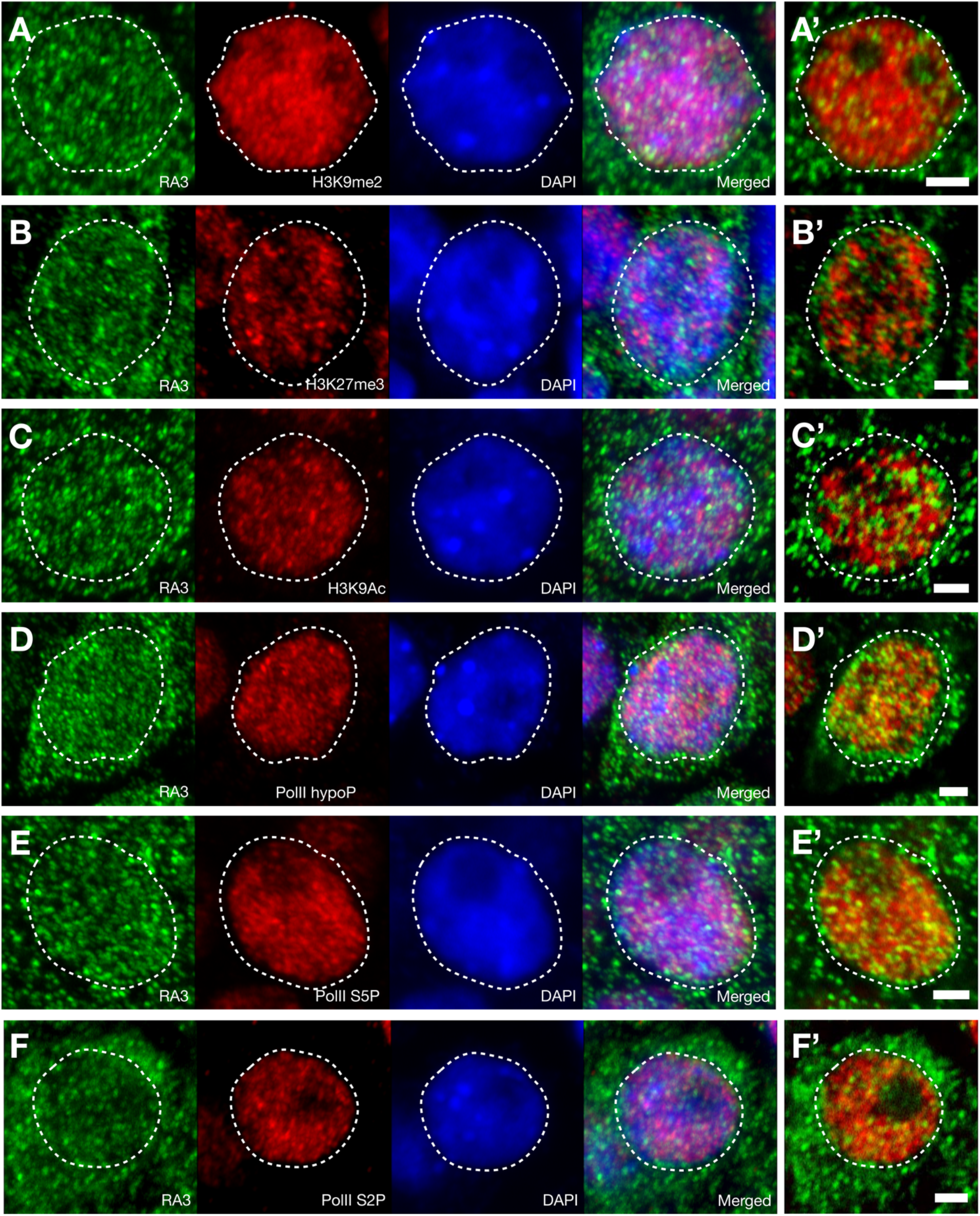
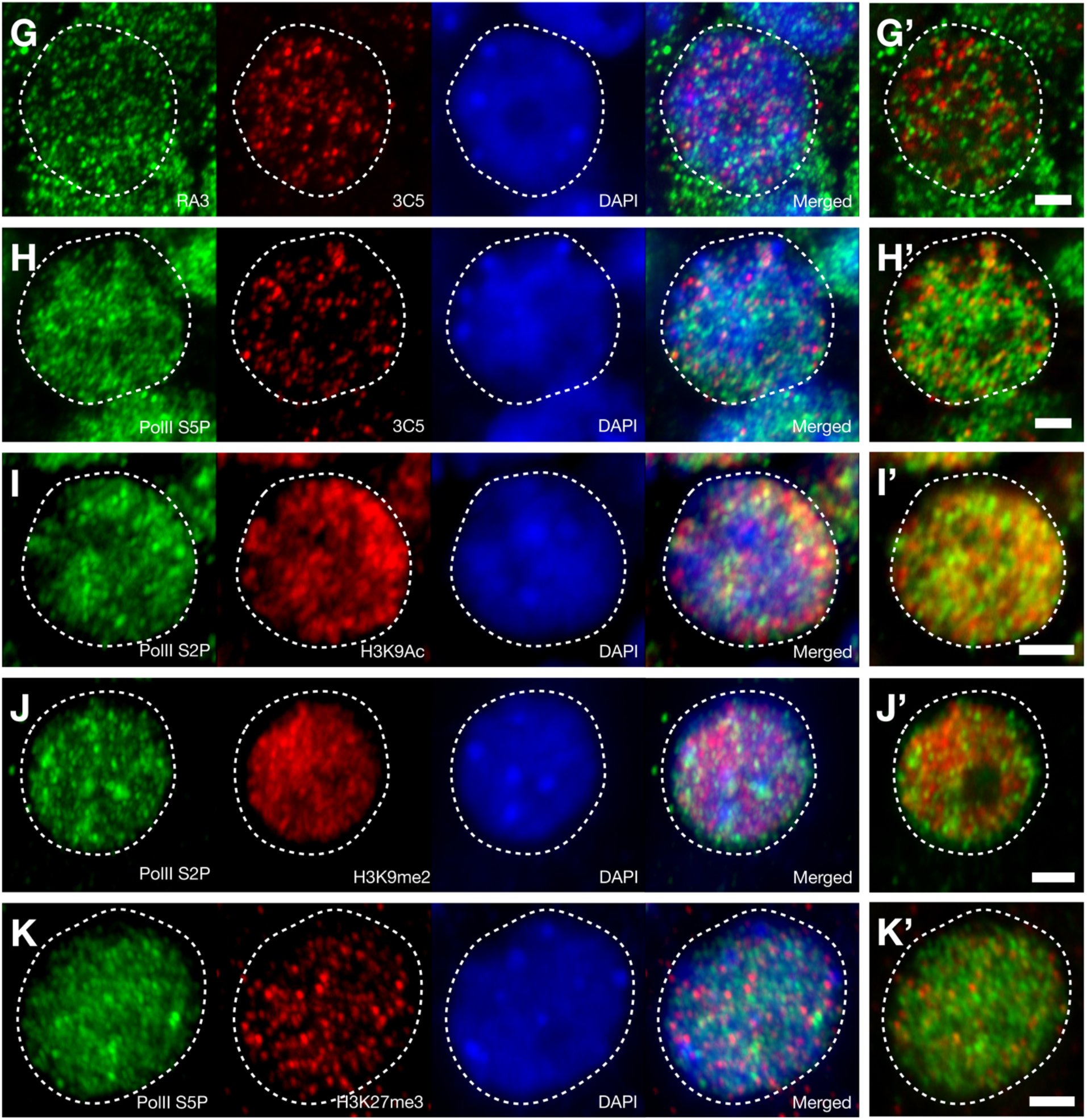
Non-processed colocalization images. Representative images of images used to estimate the 3D colocalization between RA3 (green) and nuclear markers (red, **A-G**) and between nuclear markers used as a control (**H-K**), DAPI (blue) was used for nuclear counterstaining. The images show Z-Stack projections encompassing nuclei except in **A’** to **K’** which illustrate the double colocalization in a single optical slice in the medial region of each nuclei. Scale bars: 2 μm.

**Supplemental Figure 3.**
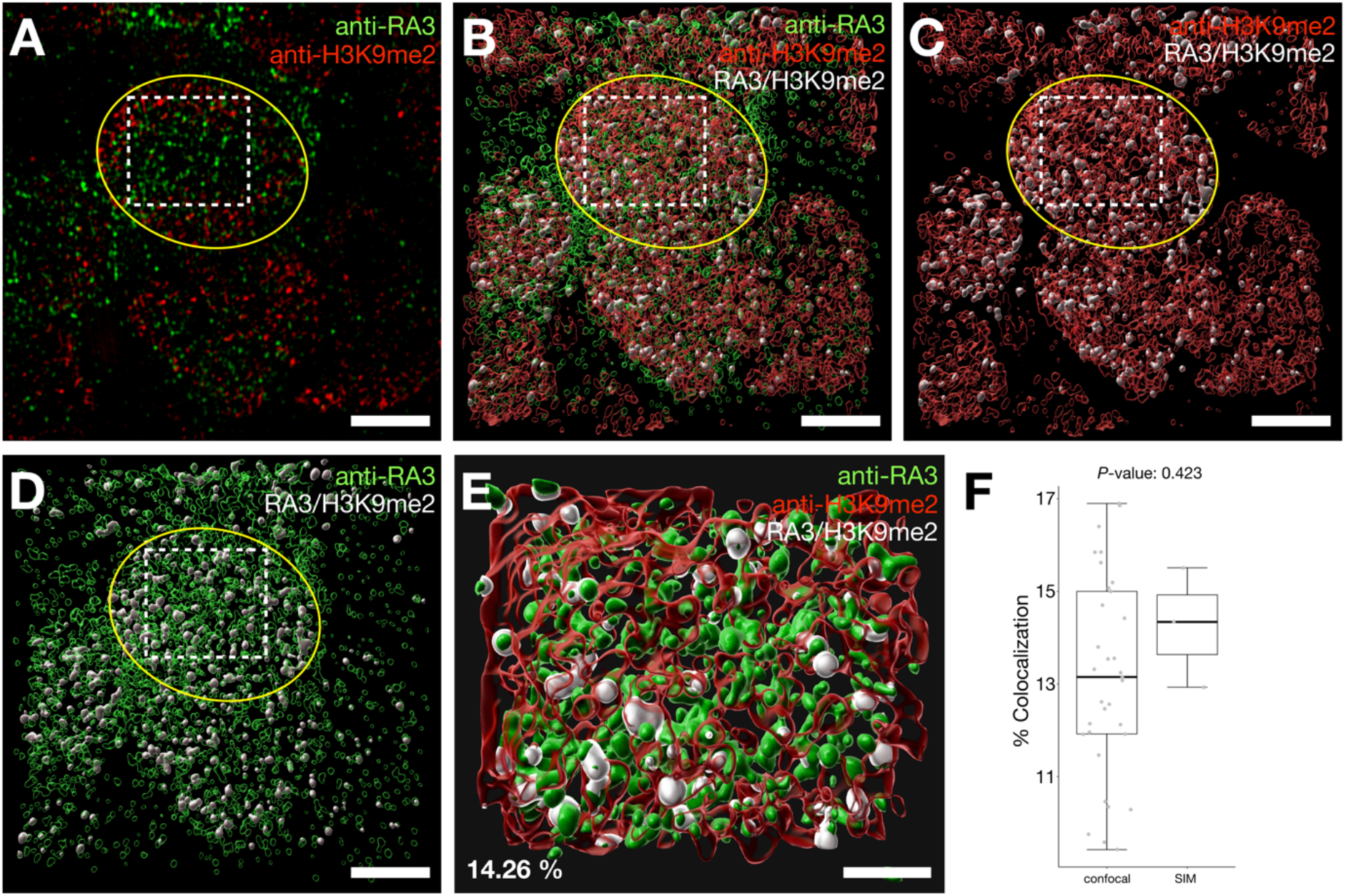
Super-resolution microscopy reveals similar 3D colocalization compared with our confocal-based approach. **A)** Representative single optical section using super-resolution structured illumination microscopy in double labelling immunolocalization using antibodies against RA3 (green) and anti-H3K9me2 (red), the nucleus is highlighted (yellow line) and a region of interest inside of the nucleus (dashed line); Z-stacks covering entire nuclei were used to quantify the volume of colocalization between RA3 and this histone mark. **B)** 3D reconstruction using the Z-stack from **A**, showing the regions where RA3 colocalized with H3K9me2 (white). **C)** Rendering showing only H3K9me2 (red) and the speckles where it colocalized with RA3 (white). **D)** RA3 (green) is localized in cytoplasm and nucleus, the regions of colocalization with H3K9me2 are showed in white. **E)** Representative region of interest inside of the nucleus, RA3 is colocalized with H3K9me2 in 14.3% of the volume. **F)** the percentage of colocalization using images acquired by super-resolution microscopy (n=3) were not significantly different compared with those acquired by confocal microscopy (n=33), the *P*-value was calculated by Mann-Whitney test. Scale bars, **A-D**: 3 μm, **E**: 1 μm.

**Supplemental Figure 4.**
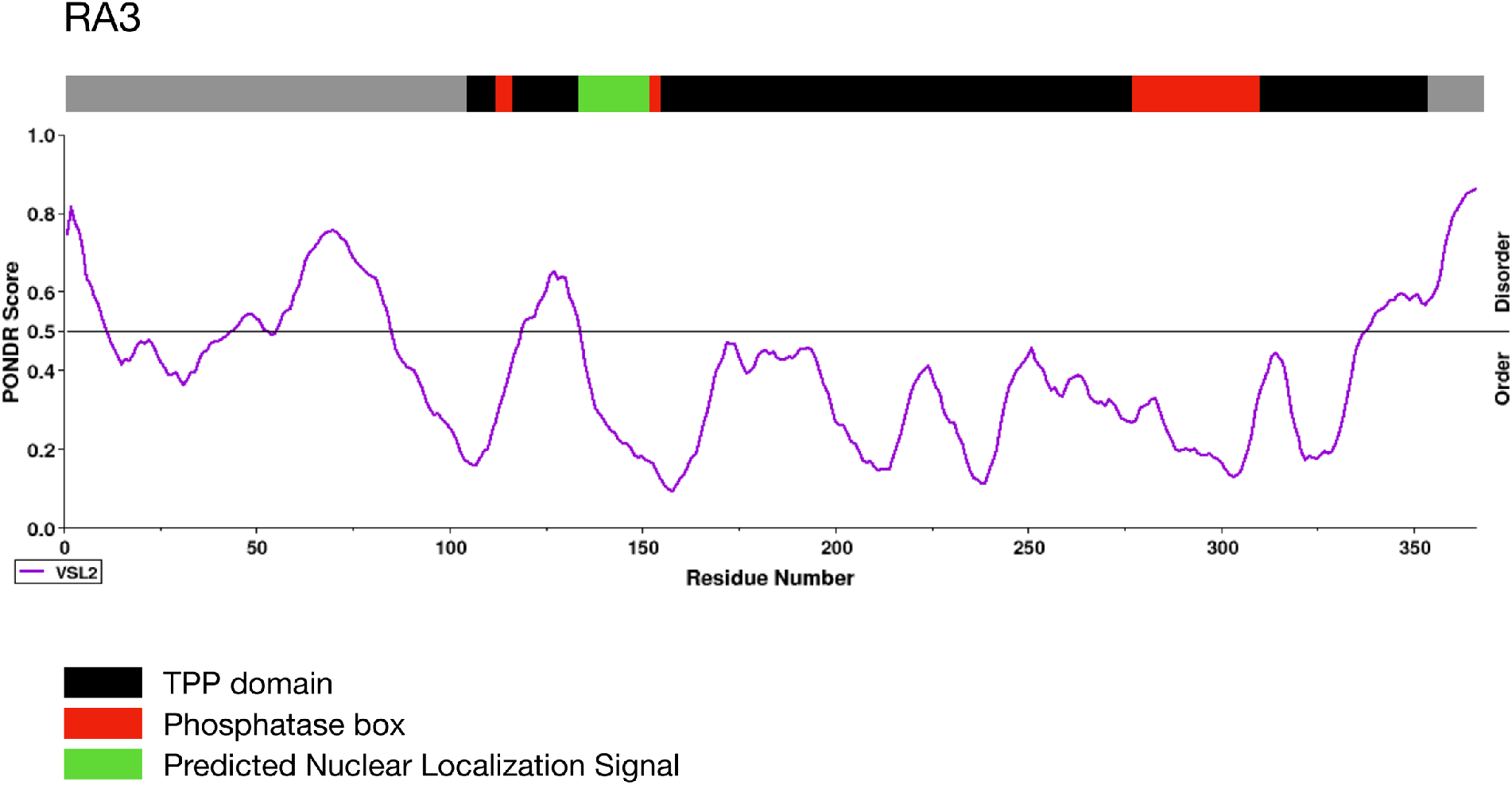
RA3 is predicted to contain a Nuclear Localization Signal and disordered regions. Prediction using LOCALIZER (http://localizer.csiro.au/) shows a Nuclear Localization Signal (green box) close to one of the phosphatase boxes (red boxes) in its TPP domain (black box). An analysis of disordered amino acid regions using PONDR (http://www.pondr.com/) shows RA3 has four putative disordered regions.

**Supplemental Table 1.** Antibodies used in this study.

| **Antibody** | **Catalog Number** | **Type** | **Supplier** | **Host** | **Dilution** |
| --- | --- | --- | --- | --- | --- |
| Anti-RAMOSA3 | NA | Polyclonal | N/A | Rabbit | 1:200 |
| Anti-H3K9me2 | C15200154 | Monoclonal | Diagenode | Mouse | 1:200 |
| Anti-H3K27me3 | C15200181 | Monoclonal | Diagenode | Mouse | 1:200 |
| Anti-H3K9Ac | C15200185 | Monoclonal | Diagenode | Mouse | 1:200 |
| Anti-H3K9me2 | C15410060 | Polyclonal | Diagenode | Rabbit | 1:200 |
| Anti-H3K9Ac | 07-352 | Polyclonal | Millipore | Rabbit | 1:200 |
| Anti-YSPTSPS repeat (RNA Pol II, B1 subunit) | C15200004 | Monoclonal | Diagenode | Mouse | 1:200 |
| Anti-YSPTSPS repeat S2Pho (RNA Pol II, B1 subunit) | C15200005 | Monoclonal | Diagenode | Mouse | 1:200 |
| Anti-YSPTSPS repeat S5Pho (RNA Pol II, B1 subunit), recombinant [EP1510Y] | ab76292 | Monoclonal | Abcam | Rabbit | 1:200 |
| Anti-YSPTSPS repeat S5Pho (RNA Pol II, B1 subunit) [H5] | ab24758 | Monoclonal | Abcam | Mouse | 1:200 |
| Anti-YSPTSPS repeat S2Pho (RNA Pol II, B1 subunit) [4H8] | ab5408 | Monoclonal | Abcam | Mouse | 1:200 |
| Anti-3C5 | NA | Monoclonal | N/A | Mouse | 1:75 |
| Anti-H3K9Ac [AH3-120] | ab12179 | Monoclonal | Abcam | Mouse | 1:200 |
| Anti-H3K9me2 [mAbcam1220] | ab1220 | Monoclonal | Abcam | Mouse | 1:200 |
| Anti-Nuclear Pore Complex proteins, clone MAb414 | 902907 | Monoclonal | BioLegend | Mouse | 1:100 |
| Anti-Mouse IgG (H+L) Highly Cross-Absorbed, Alexa Fluor 568 | A-11031 | Polyclonal | ThermoFisher | Goat | 1:300 |
| Anti-Rabbit IgG (H+L) Highly Cross-Absorbed, Alexa Fluor 488 | A-11034 | Polyclonal | ThermoFisher | Goat | 1:300 |
| Anti-Mouse IgG (H+L) Highly Cross-Absorbed, Alexa Fluor 488 | A-11029 | Polyclonal | ThermoFisher | Goat | 1:300 |
| Anti-Rabbit IgG (H+L) Highly Cross-Absorbed, Alexa Fluor 594 | A-11037 | Polyclonal | ThermoFisher | Goat | 1:300 |

